# Discovery of ester-linked ubiquitylation of PARP10 mono-ADP-ribosylation in cells: a dual post-translational modification on Glu/Asp side chains

**DOI:** 10.1101/2024.06.27.600929

**Authors:** Daniel S Bejan, Rachel E Lacoursiere, Jonathan N Pruneda, Michael S Cohen

**Affiliations:** Department of Chemical Physiology and Biochemistry, Oregon Health & Science University, Portland, OR, 97239, USA; Department of Molecular Microbiology and Immunology, Oregon Health & Science University, Portland, OR, 97239, USA

## Abstract

The prevailing view on post-translational modifications (PTMs) is that amino acid side chains in proteins are modified with a single PTM at any given time. However, a growing body of work has demonstrated crosstalk between different PTMs, some occurring on the same residue. Such interplay is seen with ADP-ribosylation and ubiquitylation, where specialized E3 ligases ubiquitylate targets for proteasomal degradation in an ADP-ribosylation-dependent manner. More recently, the DELTEX family of E3 ligases was reported to catalyze ubiquitylation of the 3’- hydroxy group of the adenine-proximal ribose of free NAD^+^ and ADP-ribose *in vitro*, generating a non-canonical ubiquitin ester-linked species. In this report, we show, for the first time, that this dual PTM occurs in cells on mono-ADP-ribosylated (MARylated) PARP10 on Glu/Asp sites to form a MAR ubiquitin ester (MARUbe). We term this process <u>m</u>ono-<u>A</u>DP-ribosyl <u>ub</u>iquit<u>ylation</u> or MARUbylation. Using chemical and enzymatic treatments, including a newly characterized bacterial deubiquitinase with esterase-specific activity, we discovered that PARP10 MARUbylation is extended with K11-linked polyubiquitin chains. Finally, mechanistic studies using proteasomal and ubiquitin-activating enzyme inhibitors demonstrated that PARP10 MARUbylation leads to its proteasomal degradation, providing a functional role for this new PTM in regulating protein turnover.

## Introduction

Post-translational modifications (PTMs) impact protein function, localization, stability, and interactions, ultimately governing key cellular processes essential for life. PTMs can range from minor alterations, like adding a methyl group (0.015 kDa), to large changes, such as conjugation to ubiquitin (∼8.5 kDa). PTMs provide proteins with a wide range of functional and structural diversity.

These modifications can happen individually, at various sites within a protein, or sometimes, multiple modifications can occur at one specific site. Two enzyme families exemplify the latter, namely PARPs and ubiquitin ligases. PARPs, a family of 17 members in humans, transfer ADP-ribose (ADPr) from nicotinamide adenine dinucleotide (NAD^+^) onto target substrates in a process called ADP-ribosylation. Mono-ADP-ribosylation (MARylation) is the transfer of a single unit of ADPr, whereas poly-ADP-ribosylation (PARylation) involves the transfer of multiple units of ADPr (up to 190 *in vitro*^1^). The PARP members are divided based on their MARylation (PARP3, 4, 6-12, 14-16) or PARylation (PARP1, 2, TNKS1, TNKS2) activity, which can occur on nucleophilic amino acids, including glutamate, aspartate, serine, arginine, and cysteine^2^. Ubiquitylation (or ubiquitination) is a multi-step process involving ubiquitin (Ub) activation by E1 enzymes, conjugation by E2 enzymes, and finally transfer onto target substrates by E3 Ub ligases. Ub is canonically attached to lysine residues of a target protein via its carboxy-terminal glycine, generating an isopeptide linkage. This monoUb species can then be extended into polyUb chains through seven different lysines (K6, K11, K27, K29, K33, K48, K63) or the amino-terminal methionine (M1) of Ub^3^.

A growing body of work indicates the existence of crosstalk between ADP-ribosylation and ubiquitylation in both prokaryotes and eukaryotes. Prokaryotes do not encode a canonical Ub system. However, bacterial pathogens can disrupt host ubiquitylation pathways by secreting specialized effector proteins^4,5^. For example, *Legionella pneumophila* secretes SdeA, a Ub ligase of the SidE family that ubiquitylates host substrates independent of ATP and canonical E1 and E2 enzymes^6^. This is achieved through concerted activities of the mono-ADP-ribosyltransferase (mART) and phosphodiesterase (PDE) domains in SdeA. First, SdeA uses its mART domain to bind NAD^+^ and MARylate Ub on Arg42. The ADPr pyrophosphate bond is then cleaved by the PDE domain, and the resulting phosphoribosyl-Ub intermediate is transferred onto a substrate serine^7^. *Chromobacterium violaceum* also secretes an effector protein, CteC, that MARylates Ub on Thr66^8^. In both cases, modification of Ub prevents activation and conjugation by E1 and E2 enzymes, respectively, resulting in significant impairments to host Ub signaling and many Ub-dependent cellular processes.

Eukaryotes also demonstrate an intricate relationship between ADP-ribosylation and ubiquitylation. A notable example is with RNF146, an E3 ligase belonging to the really interesting new gene (RING)-type class of E3 ligases, which catalyzes ubiquitylation of substrates in a PARylation-dependent manner, ultimately leading to their degradation by the Ub proteasome system^9,10^. Upon binding of iso-ADPr, an internal repeating unit of poly-ADP-ribose (PAR), to the Trp-Trp-Glu (WWE) domain of RNF146, an allosteric switch occurs that activates RNF146 ligase activity^11^. A similar finding has been observed with E3 ligases containing PAR-binding zinc finger (PBZ) motifs, namely CHFR^12,13^ and RNF114^14^, which have been shown to ubiquitylate PARylated substrates (including PARP1) for degradation. Finally, DELTEX (DTX) E3 ligases highlight an even greater interplay between ubiquitylation and ADP-ribosylation. DTX family members (DTX1, 2, 3, 3L, 4) contain a DTX C-terminal (DTC) domain that has been shown to bind PAR and mediate PAR-dependent ubiquitylation of substrates^15^. Conversely, the carboxy-terminus of Ub is MARylated upon heterodimerization of DTX3L and PARP9^16^. It was later shown that this Ub modification occurred with DTX3L alone, independent of PARP9^17^, making the role of ADP-ribosylation unclear in this model. Recently, Zhu et al. proposed an intriguing model whereby DTX E3 ligases ubiquitylate the 3’-hydroxyl of the adenine-proximal ribose of substrates, including free NAD^+^ or ADPr^18^, and even ADP-ribosylated nucleic acids^19^. In this model, ADPr is ubiquitylated rather than Ub being ADP-ribosylated.

Using in vitro systems, these studies suggest an exciting and novel way for ADPr and Ub to occur at the same site on proteins as a dual PTM. However, a crucial question remains whether this dual post-translational modification occurs on proteins within cells, and if it does, what significance it holds in terms of function. Herein, we show for the first time that MARylated PARP10 is polyubiquitylated in cells on a hydroxyl group within the adenine-proximal ribose (A-ribose) of MARylated PARP10, generating a noncanonical ester-linked mono-ADPr Ub species (MARUbe). Moreover, we demonstrate that on PARP10, this dual PTM, which we call <u>m</u>ono-<u>A</u>DP-ribosyl <u>ub</u>iquit<u>ylation (MARUbylation)</u>, is extended with K11-linked polyUb chains, which we suggest is important for regulating PARP10 turnover by the Ub proteasome system.

## Results

### PARP10 high molecular weight MARylation is dependent on ubiquitin

In 2008, PARP10 (formerly known as ARTD10^20^) was the first member of the PARP family shown to possess MARylating activity^21^. Although when PARP10 was first discovered three years prior, it was originally proposed to possess auto-PARylating activity^22^, as evidenced by weak detection with a monoclonal antibody specific for poly-ADPr^22,23^. These experiments were done *in vitro*, so it was still unclear what kind of activity PARP10 displayed in cells, especially since MARylation had not been detected then. This was largely due to the lack of antibodies/detection reagents specifically recognizing mono-ADPr versus poly-ADPr. Since then, new reagents specific for mono-ADPr^24–26^ and pan mono/poly-ADPr reagents^24,27^, show that PARP10 is auto-MARylated in cells^28^. In fact, most of the PARP family members display MARylation activity in cells^28^. It was therefore striking to us that following expression of PARP10 using doxycycline (dox)-inducible HEK 293 cells, we observed an upward mobility shift (or “smear”) of ADP-ribosylation originating at the molecular weight of GFP-PARP10, detected using a mono/poly-ADPr antibody (**Figure 1A**). This led us to conclude that, in our system, the high molecular weight (MW) PARP10-mediated ADP-ribosylation smear is evidence of auto-PARylation of PARP10. However, the high MW smear of ADP-ribosylation was also detected with a mono-ADPr-specific antibody^25^, but not with a poly-ADPr-specific detection reagent^24,25^, indicating the absence of any PARylation (**Figure 1A**). To confirm these antibodies/detection reagents are working as expected, we transiently overexpressed GFP-PARP10 in HEK 293 control and PARP1 KO cells to generate a MARylation signal or treated with a poly(ADP-ribose) glycohydrolase (PARG) inhibitor^29^ to stabilize an endogenous PARP1-mediated PARylation signal (**Figure S1**). As expected, we detected PARP1-dependent PARylation with the poly- and mono/poly-ADPr antibodies, but not with the mono-ADPr-specific antibody. Conversely, GFP- PARP10 MARylation was detected with the mono- and mono/poly-ADPr antibodies, but not with the poly-ADPr-specific antibody (**Figure S1**).

**Figure 1.**
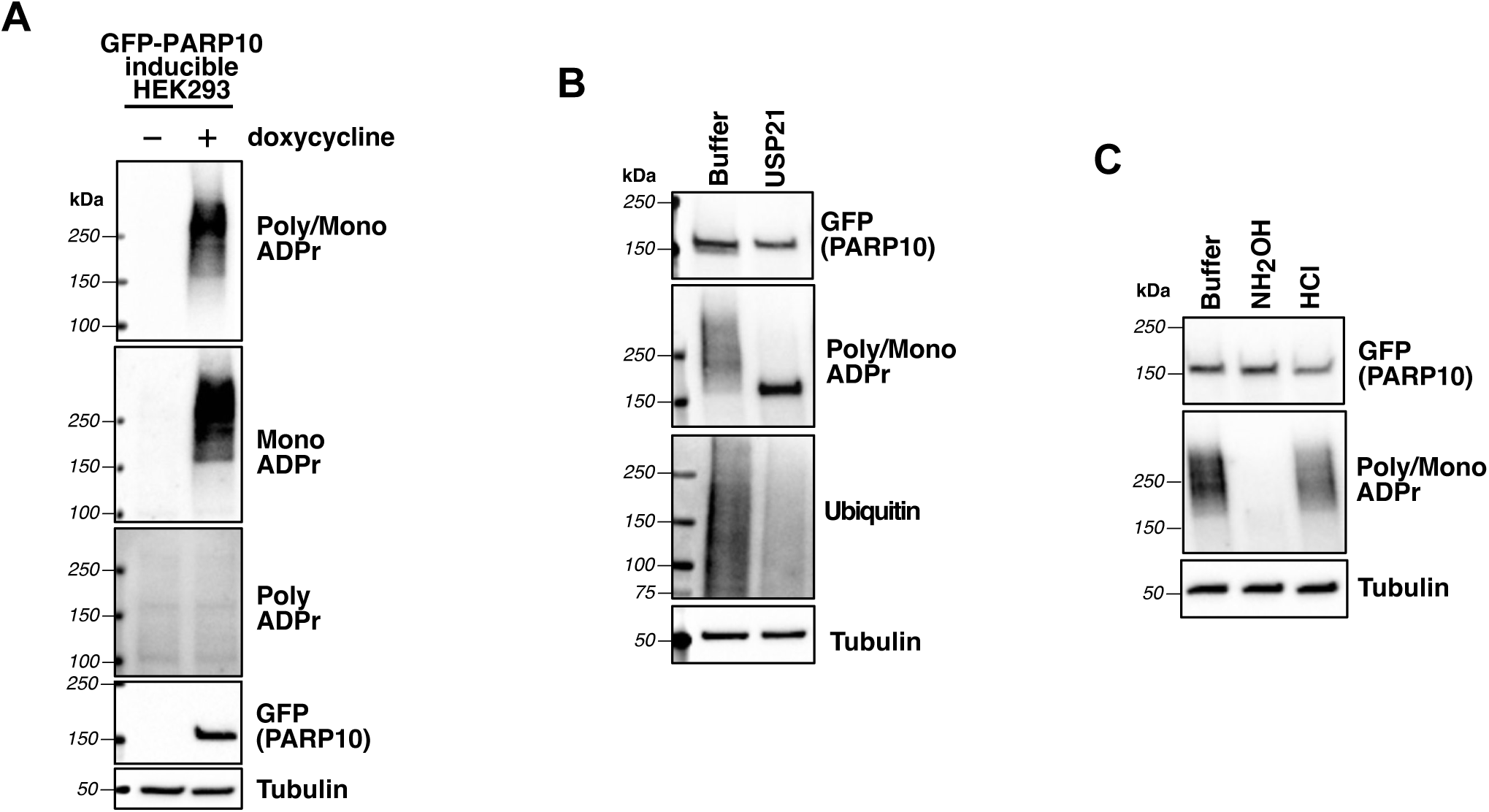
High molecular weight PARP10 auto-MARylation is ubiquitin dependent. (A) HEK 293 GFP-PARP10 doxycycline-inducible cells were treated with doxycycline (10 µg/ml) for 24 hours, followed by western blotting and probing for poly/mono ADPr (Cell Signaling Technology: E6F6A), mono-ADPr (Bio-Rad: HCA354), or poly-ADPr (Millipore Sigma: MABE1031). (B) HEK 293 cell lysates containing doxycycline-induced GFP-PARP10 were treated with 1 µM USP21 for 1 hr at 37°C, followed by western botting. (C) HEK 293 cell lysates containing doxycycline-induced GFP-PARP10 were treated with hydroxylamine (1M, pH 7) or HCl (300 mM, pH 1.5) for 15 minutes, followed by methanol precipitation and western blotting (HCl-treated samples were neutralized with equimolar NaOH prior to precipitation).

What could be underlying the high MW PARP10 MARylation smear, given that PARylation is not involved? PARP10 is the only PARP that contains Ub-interacting motifs (UIMs)^30^, which have been shown *in vitro* to bind K63-linked tetraUb^31^ and K48-linked diUb^32^, independently. PARP10 has also been shown to be polyubiquitylated by RNF114 in cells^33^. Therefore, we hypothesized that the high MW PARP10 MARylation smear could be due to polyubiquitylation. To test this idea, we incubated lysates containing dox-induced GFP-PARP10 with USP21, a nonspecific deubiquitinase (DUB) that removes all polyUb linkage types^34^. Upon USP21 treatment, we observed a collapse of the high MW MARylated smear into a distinct mono-ADPr band at the molecular weight of GFP-PARP10 (**Figure 1B**), indicating that polyubiquitylation is responsible for the high MW PARP10 MARylation smear. We also wanted to determine which amino acid(s) on PARP10 are MARylated. Previous studies^21,35–37^ demonstrated acidic amino acids, such as glutamate (Glu) and aspartate (Asp), are MARylated on PARP10. Therefore, we treated lysates with neutral hydroxylamine, a chemical that can rapidly cleave ester-linked ADPr^38,39^ and observed the removal of the MARylation smear (**Figure 1C**). As a negative control, we subjected lysates to 300 mM HCl (pH 1.5) and observed no removal of MARylation, indicating that Glu/Asp-linked ADPr is stable in acidic conditions, as previously reported^40^. These results support the hypothesis that the high MW PARP10 MARylation smear is due to polyubiquitylation of PARP10 and that PARP10 MARylation occurs exclusively on Glu/Asp side chains in cells.

### An engineered bacterial DUB with profound Ub esterase activity cleaves a Ub ester-linked to ADPr on a model substrate

There are two major ways proteins can be modified with Ub. One is through a lysine, which occurs via a canonical isopeptide linkage. The other is ubiquitylation of serine or threonine, generating an ester-linkage (or with a cysteine, leading to a thioester). Ester-linked Ub may also be conjugated to proteins through ubiquitylation of the covalently attached ADPr group, as previously suggested *in vitro*^18^. While hydroxylamine has previously been used to diagnostically release ester-linked ubiquitylation^41–43^, in our case, its use would be confounded by our finding that PARP10 is Glu/Asp-MARylated in cells. Recently, the bacterial deubiquitinase (DUB) TssM from *Burkholderia pseudomallei* was shown to reverse ester-linked lipopolysaccharide ubiquitylation as a strategy to evade host cell-intrinsic immune responses^44^. Using Lys-Ub and Ser/Thr-Ub models of isopeptide- and ester-linked Ub, respectively, TssM was 30 times more active towards ester-linked Ub than isopeptide-linked Ub. Guided by a Ub-bound crystal structure of TssM, a V466R variant (referred to here as TssM*) was identified that further uncoupled esterase and isopeptidase activities, enhancing its esterase-specific activity by ∼5,000-fold^44^. We, therefore, reasoned that TssM and TssM* could potentially be used to determine if PARP10 is ubiquitylated on Glu/Asp-MARylation sites in cells.

Before testing TssM and TssM* on MARylated PARP10 isolated from our dox-inducible cells, we wanted to validate their specificities against a model substrate containing an ester-linked Ub at the 3’-OH of the proximal adenine-ribose (A-ribose) of NAD^+^. Following Zhu et al^18^, we used DTX2 to ubiquitylate an NAD^+^ analog containing a desthiobiotin (DTB) at the C2 position of the adenine ring (DTB-NAD^+^) ^45,46^ (**Figure 2A**). We found that DTX2 could readily conjugate Ub to the 3’-OH of the A-ribose of DTB-NAD^+^ to form Ub ester-linked DTB-NAD^+^ (DTB-NAD^+^-Ube), as evidenced by the presence of a band ∼10 kDa (detected using streptavidin-HRP), which is the expected molecular weight of DTB-NAD^+^-Ube (**Figure 2B**). We found that TssM and TssM* efficiently cleaved the Ub ester at the 3’-OH of the A-ribose of DTB-NAD^+^ as evidenced by the disappearance of the ∼10 kDa DTB-NAD^+^-Ube band in the streptavidin-HRP blot (**Figure 2B**). As an orthogonal strategy, we showed hydroxylamine also eliminated the band corresponding to DTB-NAD^+^-Ube. In addition to ester-linked ubiquitylation of NAD^+^, DTX2 was also reported to catalyzes isopeptide-linked auto-ubiquitylation in the same reaction^18^. We detected this activity as well, which served as a convenient internal control for DUB specificity. As expected, TssM, but not TssM* or hydroxylamine, cleaved DTX2 auto-ubiquitylation (**Figure 2B**), suggesting DTX2 auto-ubiquitylation is isopeptide-linked and that TssM* is indeed a useful tool for ester-specific Ub removal. Finally, we used maltoheptaose-Ub, another sugar-containing ester-linked Ub substrate, to demonstrate the esterase-specific activity of TssM*. Maltoheptaose is ubiquitylated on a C6 hydroxyl group of a glucosyl unit (**Figure S2A**) by the E3 ligase HOIL-1 in the presence of K63-linked diUb^47^. Similar to DTB-NAD^+^-Ube, the maltoheptaose-Ub signal was removed by TssM, TssM*, and hydroxylamine. The isopeptide-linked K63 diUb substrate (serving as an internal specificity control), however, was only cleaved by TssM (**Figure S2B**). Taken together, these results confirm that TssM is an active DUB with dual esterase and isopeptidase activity, and that the TssM* variant displays exquisite esterase-specific DUB activity for Ub attached to hydroxyl groups on different types of sugars.

**Figure 2.**
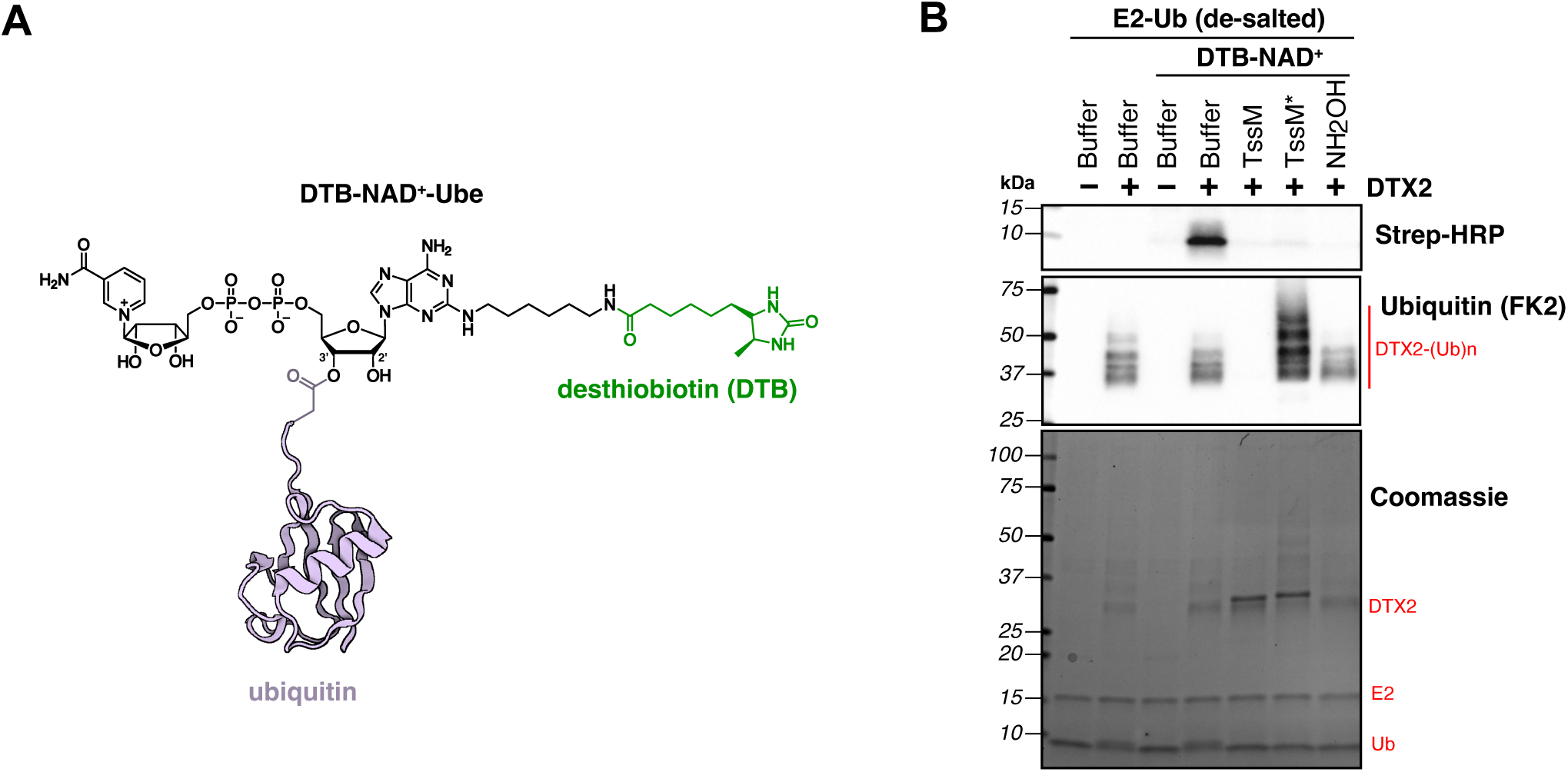
TssM* is highly specific for removal of ester-linked Ub on sugars. (A) Structure of DTB-NAD^+^ with ester-linked Ub on the 3’-OH group of ribose (DTB-NAD^+^-Ube). (B) DTX2 was incubated with 25 µM DTB-NAD^+^ and a loaded E2-Ub complex (de-salted) for 1 hr at 37°C, followed by 1 hr reaction at 37°C with 1 µM TssM, 1 µM TssM*, or 1M NH_2_OH (pH 7.5).

### Enzymatic and chemical sensitivity studies reveal that PARP10 is ubiquitylated on MARylated Glu/Asp sites in cells

Having validated TssM* as an esterase-specific DUB capable of cleaving Ub ester-linked to ADPr, we asked if PARP10 is ubiquitylated at the 3’-OH group of the A-ribose of Glu/Asp- MARylation sites. We transiently expressed tagged Ub (HA-Ub) in our GFP-PARP10 dox-inducible cells and immunoprecipitated GFP-PARP10 using GFP-Trap beads. After stringent washes (with 7M urea and 1% SDS) to remove noncovalently interacting ubiquitylated proteins, we performed on-bead enzymatic or chemical treatments (**Figure 3A**), followed by western blotting to detect changes in MARylation and ubiquitylation on PARP10 (**Figure 3B**). Treatment with TssM removed nearly all Ub from PARP10, resulting in a collapse of the high MW ADPr smear to a single band at the MW of GFP-PARP10, as seen with USP21 treatment (**Figure 1C**). Remarkably, treatment with TssM* removed ∼40% of the Ub from PARP10, partially collapsing the high MW ADPr smear into a single band (**Figure 3B, 3C**). These results show that a significant population of Ub on PARP10 is ester-linked in cells.

**Figure 3.**
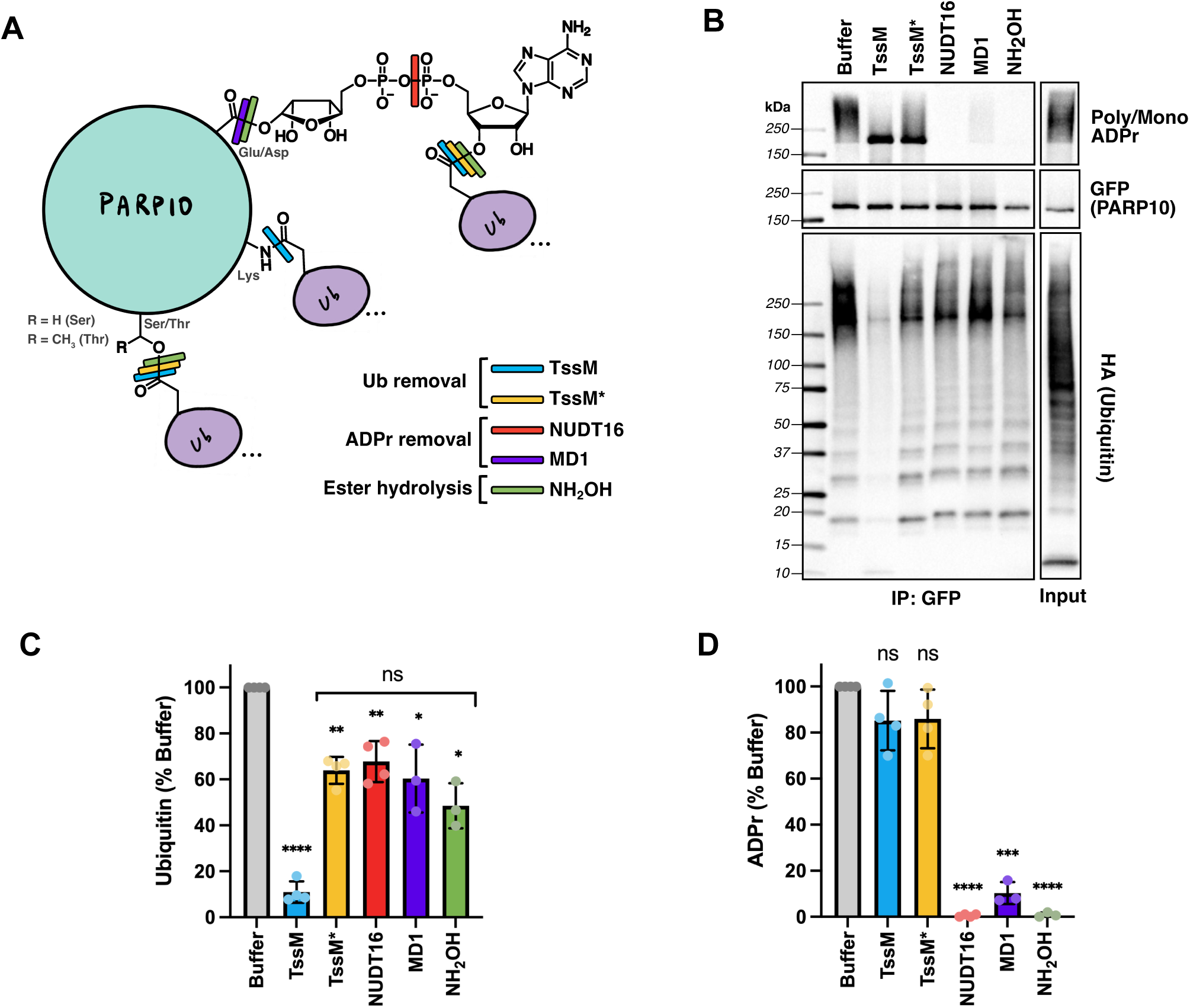
Polyubiquitin is directly attached to a hydroxyl group on ADP-ribose of MARylated PARP10 in cells. (A) Schematic of potential sites for Ub attachment on MARylated PARP10, with corresponding cut sites for different enzymatic or chemical treatments. (B) GFP-PARP10 from doxycycline- treated HEK 293 cells was immunoprecipitated with GFP-trap beads, washed stringently (7M Urea, 1% SDS), and beads were treated with DUBs, ADPr hydrolases, or chemicals, followed by western blotting. Quantification of the ∼150 kDa ubiquitin smear (C) and ADPr smear (D) from (B); n = 3-4 biological replicates. *p<0.05, **p<0.01, ***p<0.001, ****p<0.0001, one sample t and Wilcoxon test relative to buffer treatment. ns = not significant. For (C), comparison between TssM*, NUDT16, MD1, and NH_2_OH was measured by one-way ANOVA (ns).

Ester-linked Ub could be on Ser/Thr residues or the 3’-OH of the attached ADPr (**Figure 3A**). To distinguish these possibilities, we treated samples with Nudix-Type Motif 16 (NUDT16), a hydrolase that cleaves MARylation and PARylation at the pyrophosphate bond in ADPr^48^ (**Figure 3A**). If NUDT16 decreases the high MW PARP10 Ub smear, this would support the hypothesis that Ub is ester-linked to the 3’-OH of the A-ribose of ADPr. If no change in PARP10 ubiquitylation is observed with NUDT16 treatment, this would suggest that Ub is ester-linked to a Ser/Thr on PARP10. Upon treatment with NUDT16, we observed complete removal of MARylation (**Figure 3B, 3D**) and a ∼40% reduction in Ub, similar to TssM* treatment (**Figure 3B, 3C**), suggesting that Ub is indeed linked to the 3’-OH of the A-ribose of ADPr. Furthermore, we found that treatment with hydroxylamine or MacroD1 (MD1), an ADPr hydrolase that cleaves Glu/Asp-linked ADPr^49,50^, also resulted in a ∼40% reduction in Ub (**Figure 3C**), with near complete removal of MARylation (**Figure 3D**). We also tested other macrodomain-containing ADPr hydrolases (MacroD2 (MD2) and terminal ADP-Ribose protein glycohydrolase (TARG)^51^), and observed a similar effect, with TARG displaying more potent removal of PARP10 MARylation and subsequent Ub release (**Figure S3**). Together, these results demonstrate that PARP10 is polyubiquitylated in cells, and nearly half of this polyUb is ester-linked to the 3’-OH of the A-ribose of Glu/Asp-MARylated PARP10. We refer to this dual modification as a mono-ADPr-Ub ester (MARUbe) and the novel PTM as <u>m</u>ono-<u>A</u>DP-ribosyl <u>ub</u>iquit<u>ylation</u>, or MARUbylation.

### PARP10 MARUbe contains K11-linked polyubiquitin

We next sought to further characterize the population of ester-linked polyUb on ADPr and the ∼60% polyUb remaining on PARP10 after TssM* or ADPr hydrolase treatment. To distinguish these two populations, we stringently washed immunoprecipitated GFP-PARP10 as described above; however, after on-bead treatment with NUDT16, we separated the supernatant (containing released polyUb from ADPr) and beads (containing polyUb retained on PARP10) (**Figure 4A**). These fractions were then treated with the nonspecific DUBs TssM and USP21 or a panel of polyUb linkage-specific DUBs in a workflow termed Ubiquitin Chain Restriction Analysis (UbiCRest)^52,53^ (**Figure 4B**). In the beads fraction, the remaining polyUb on PARP10 after NUDT16 treatment was efficiently removed by the nonspecific DUBs but not with TssM*, suggesting canonical isopeptide-linked polyUb. Intriguingly, this polyUb species was not removed by any linkage-specific DUBs, suggesting it is either highly complex or multi-monoUb modified rather than polyubiquitylated. As an alternative strategy to characterize the isopeptide-linked polyUb on PARP10 and to test if it is dependent on MARylation, we inhibited MARylation in cells before immunoprecipitation, using a pan-MARylating PARP inhibitor, RBN010860^54^. RBN010860 potently inhibited PARP10 MARylation with a half-maximal inhibitory concentration (IC_50_) of 40.9 nM (**Figure S3A, S3B**), similar to the reported K_d_ of 34 nM measured for PARP10 *in vitro*^54^. We observed that even after near complete inhibition of PARP10 MARylation using RBN010860, a polyUb signal on PARP10 was still present and was cleaved by TssM but not TssM*, suggesting an isopeptide-linked polyUb species that is not dependent on MARylation (**Figure S4**).

**Figure 4.**
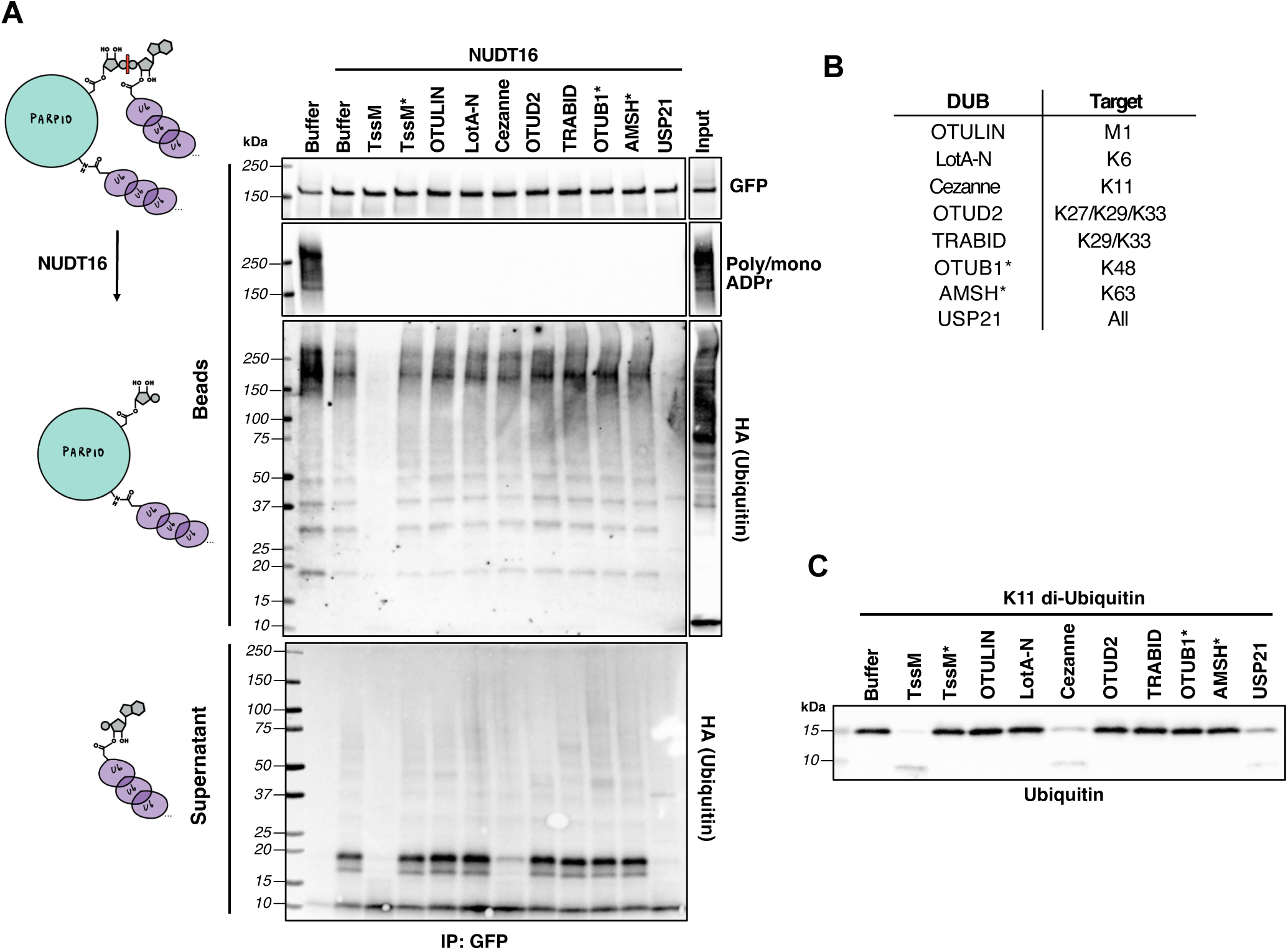
PARP10 MARUbe contains K11-linked Ub chains. (A) GFP-PARP10 from doxycycline-induced HEK 293 cells was immunoprecipitated with GFP-trap beads, washed stringently (7M Urea, 1% SDS), and beads were treated with NUDT16. The supernatant and beads were separated and further treated with polyubiquitin linkage-specific DUBs, followed by western blotting. Representative image from n = 2 biological replicates. (B) Table of polyubiquitin linkage-specific DUBs and their preferential cut site. (C) Positive control experiment with polyubiquitin linkage-specific DUBs using K11 di-ubiquitin substrate (1 µM).

In the supernatant fraction, we observed a release of polyUb upon NUDT16 treatment, with the major species being a diUb linkage. From the polyUb linkage-specific DUBs tested, only Cezanne could significantly cleave this diUb species (**Figure 4A**). Cezanne/OTUD7B, a member of the ovarian tumor (OTU) family of DUBs, was the first DUB shown to preferentially cleave K11-linked polyUb^55^. We confirmed this result using a K11-linked diUb substrate, demonstrating that Cezanne is K11-specific under our conditions (**Figure 4C**). Taken together, these results demonstrate that PARP10 contains two polyUb species; i) K11-linked polyUb attached to MARUbe and ii) canonical isopeptide-linked polyUb with an unknown, likely complex, chain identity.

### Inhibition of the proteasome or ubiquitylation system differentially impact PARP10 protein levels and MARylation

Having characterized K11-linked MARUbylation of PARP10, we next wanted to explore the functional roles of PARP10 MARUbe in cells. A common role for polyubiquitylation is to target substrates for proteasomal degradation. As discussed earlier, select E3 ligases target substrates for ubiquitin-proteasome-mediated degradation in an ADP-ribosylation-dependent manner. Therefore, we wanted to test if MARUbylation of PARP10 leads to its degradation and if this degradation is MARylation-dependent. From cell-based experiments using RBN010860, we observed that a dose-dependent decrease in PARP10 MARylation is accompanied by a dose-dependent increase in PARP10 protein levels, up to ∼2-fold (**Figure S4C**). This suggests that MARylation of PARP10 may lead to its degradation through the Ub proteasome system. To address this idea more directly, we treated dox-inducible GFP-PARP10 cells with MG132, a common inhibitor used to block the proteasome^56^ (**Figure 5A**). While levels of the unmodified GFP-PARP10 band (∼170 kDa) did not change significantly, we did observe a time-dependent stabilization of a higher MW GFP-PARP10 band (∼250 kDa) upon MG132 treatment, which correlated with a time-dependent increase in MARylation (**Figure 5A, 5B**). This supports the hypothesis that a subpopulation of PARP10, which contains mass-shifted MARylation, is degraded by the proteasome. On the other hand, we wanted to test the impact of ubiquitylation on GFP-PARP10 levels and activity. To block global ubiquitylation, we used TAK-243, a covalent E1 enzyme inhibitor that forms an adduct with the carboxy-terminus of Ub, thereby preventing the first step of Ub activation^57^. By blocking ubiquitylation of newly synthesized GFP- PARP10 molecules with TAK-243, the degradation of pre-existing MARUbylated GFP-PARP10 molecules can be monitored over time. Consistent with the model of proteasomal degradation, treatment with TAK-243 led to a time-dependent decrease in the upper GFP-PARP10 band, accompanied by a decrease in MARUbylation (**Figure 5A, 5B**). Overall, these results support a model in which PARP10 levels are regulated by K11-linked MARUbylation-dependent proteasomal degradation.

**Figure 5.**
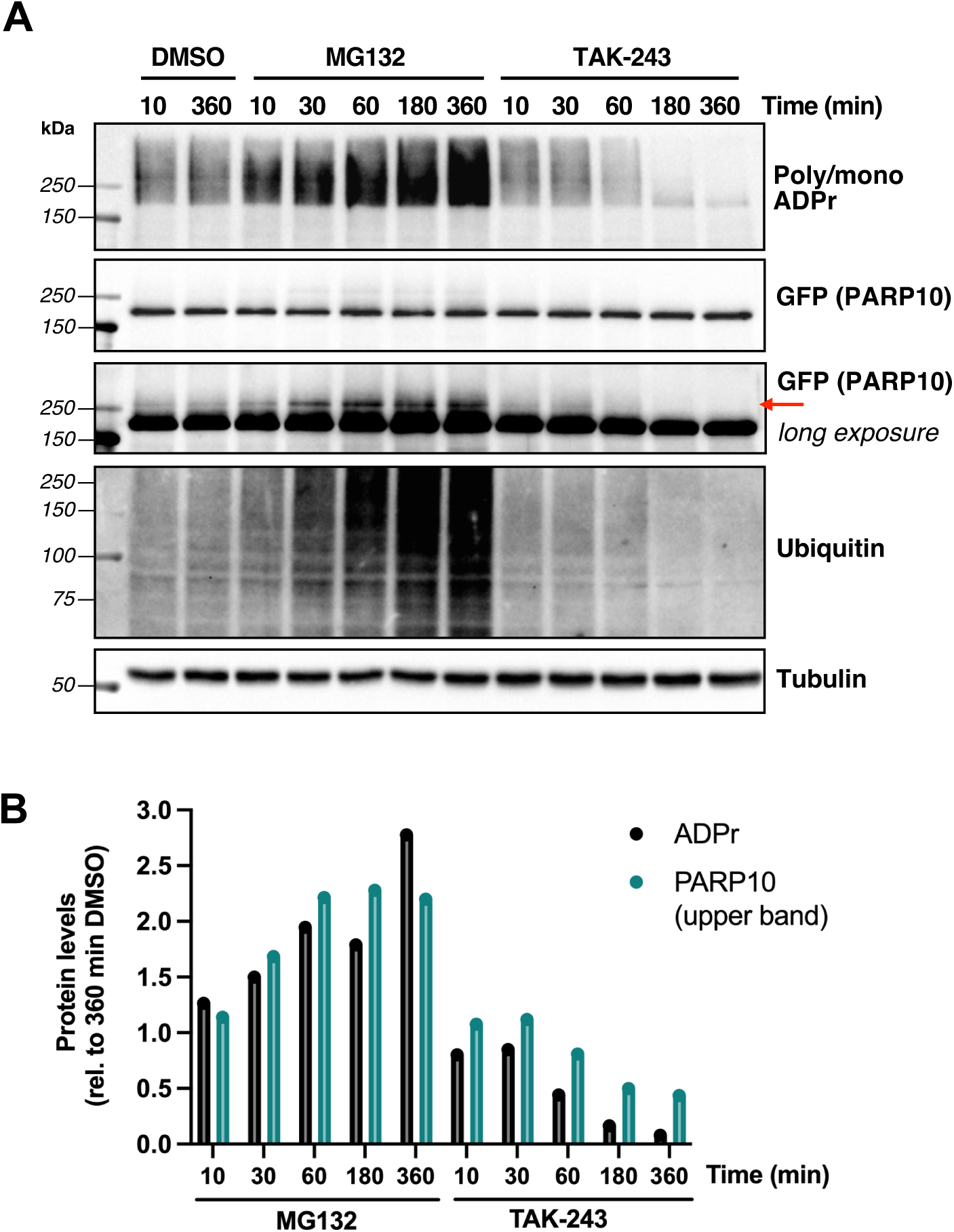
Inhibition of the proteasome or global ubiquitylation leads to opposite effects on PARP10 levels and MARylation. (A) HEK 293 GFP-PARP10 doxycycline-inducible cells were co-treated with doxycycline and MG132 (10 µM) or TAK-243 (1 µM) on a time course, followed by western blotting. (B) Quantification of the poly/mono ADPr signal and higher MW GFP-PARP10 band (indicated with a red arrow) from (A); n = 1 biological replicate.

## Discussion

The prevailing view in biology is that a PTM-targeted amino acid is modified with a single PTM (e.g., methylation of lysine, phosphorylation of serine) or, in some cases, multiple PTMs of the same type (e.g., di- and trimethylation of lysines). Intriguingly, however, there are situations where a single amino acid could be dually modified with two distinct PTMs. One study showed that a single lysine in histones can contain acetyl and methyl groups, forming a novel dual modification called acetyl-methyllysine (Kacme)^58^. Another dual-type modification is protein pyrophosphorylation, where a phosphorylated serine is phosphorylated again, generating pyrophosphoserine^59,60^. Lastly, Ub can also undergo acetylation (on lysines) or phosphorylation (on Ser/Thr/Try)^3^. In this report, we show, for the first time, that PARP10 is a MARUbylated on Glu/Asp sites in cells. This dual modification, MARUbe, contains a Ub ester conjugated to the mono-ADPr attached to Glu/Asp sites on PARP10.

We observed a greater degree of MARUbylation when PARP10 was expressed using a dox-inducible system compared to transient overexpression, which instead led to more of a single MARylated band at the molecular weight of GFP-PARP10 (**Figure 1A, S1**). One explanation for this difference could be due to the higher levels of PARP10 obtained with transfection compared to the dox-inducible system, leading to supraphysiological MARylation on non-Glu/Asp sites. Another observation we noticed is that while PARP10 auto-MARylation is highly mass-shifted (detected by poly/mono-ADPr antibody), the overall levels of PARP10 are not (detected by GFP antibody), suggesting only a small fraction of PARP10 molecules are modified in this manner (**Figure 3B**). Indeed, proteomic studies characterizing endogenous PARP1-mediated ADP-ribosylation revealed less than 11% of ADP-ribosylation occupancy in half of the ADP-ribosylated sites identified^61^. In contrast, bacterial ADP-ribosyltransferases modify their substrates much more efficiently, with some showing a 1:1 stoichiometry for target residue modification^62^. This highlights the more transient and dynamic nature of eukaryotic ADP-ribosylation compared to bacterial ADP-ribosylation, which may be governed by evolution for substrate specificity, amino acid preference, or susceptibility to ADPr hydrolases^2^. An alternative explanation for the low stoichiometry of PARP10 modification could be due to the inherent lability of esters during sample preparation. This is evident from recent reports showing that ester-linked PTMs, such as Glu/Asp-ADPr or ADPr-Ub, are highly sensitive to heat and alkaline conditions^63,64^. Therefore, standard western blotting procedures that involve boiling samples and resolving proteins in gels/buffers at pH ∼8, as performed in this study, may significantly reduce the amount of PARP10 MARUbylation detected.

Our findings suggest a role for MARUbylation in regulating PARP10 turnover through the proteasome. The typical mark for proteasomal degradation is K48-linked polyUb. While we did not detect K48-linked polyUb on PARP10 using our UbiCRest assay (**Figure 4**), this does not exclude the possibility. For example, it has been shown that branched K11/K48-linked polyUb can also mark substrates for proteasomal degradation even more efficiently than homotypic K48-linked polyUb^65,66^. Furthermore, it has been reported that Cezanne cleaves branched K11/K48-linked polyUb preferentially over homotypic K11-linked polyUb^67^. Therefore, it is possible that the K11-linked polyUb detected on the ADPr of PARP10 could also contain K48- linked Ub, and that this branched species is what targets PARP10 for rapid proteasomal degradation (**Figure 5A**). A more complex species of Ub, including branched, mixed, or multi- monoUb, could also explain why we don’t see the removal of the isopeptide-linked polyUb on PARP10 with any of the linkage-specific DUBs (**Figure 5A**). Recently, two independent proteomic studies (using orthogonal methods and distinct cell lines) exploring the interactome of branched K48/K63-linked polyUb identified PARP10 as a major target^68,69^. This finding is in line with previous studies showing that PARP10 UIMs interact independently with K63-linked^31^ and K48-linked^32^ Ub substrates *in vitro*. It is also possible that PARP10 itself is modified with branched K48/K63-linked polyUb, which can interact with PARP10 UIMs in an intramolecular fashion, thereby competing with binding to K48/K63-linked polyUb substrates. One substrate PARP10 has been proposed to interact with is polyubiquitylated TRAF6, an E3 ligase involved in the nuclear factor-κB (NF-κB) signaling pathway^31^. The NF-κB family of transcription factors regulate key processes involved in innate and adaptive immune responses^70^. Both PARP10 catalytic activity and UIMs were necessary to prevent NF-κB activation and translocation from the cytosol to the nucleus, leading to suppression of NF-κB-dependent gene and protein expression upon interleukin-1β (IL-1β) or tumor necrosis factor-α (TNFα) stimulation. The authors proposed a model where binding of PARP10 UIMs to K63-linked polyubiquitylated TRAF6 led to MARylation of NF-κB essential modulator (NEMO), ultimately leading to impaired downstream signaling and no NF-κB-dependent gene expression^31^. Moreover, it has been shown that upon IL-1β stimulation, the E3 ligase HUWE1 adds K48-linked branches to the K63- linked polyUb chains generated by TRAF6. This protects the K63 chains from K63-targeted DUBs, resulting in amplified NF-κB signaling^71^. This study further supports the hypothesis that PARP10 acts as a negative regulator of NF-κB signaling by interacting with TRAF6-branched K48/K63-linked polyUb through its UIMs.

Our study uncovered a new dual modification, MARUbe, on PARP10 in cells. MARUbe consists of K11-linked ubiquitin, which we propose regulates PARP10 stability in cells. This adds to the growing crosstalk between ADP-ribosylation and ubiquitylation, highlighting the remarkable complexity of PTMs and their influence on signaling pathways. While there are still many unanswered questions, we can say that *all good things come in pairs*…of esters; one for ADP-ribosylation and one for ubiquitylation.

## Acknowledgements

We thank current and former members of the Cohen and Pruneda lab for many insightful discussions relating to experimental design, data analysis and interpretation, and general advice and feedback for this project. We thank Dr. Sunil Sundalam (former member of Cohen lab) for synthesizing RBN010860 and Dr. Maike Lehner (former member of Prof. Andreas Marx, University of Konstanz) for synthesizing DTB-NAD^+^. This work was supported by the National Institutes of Neurological Disorders and Stroke 2R01NS088629 (to M.S.C.) and by the National Institute of General Medical Sciences funding grant R35GM142486 (to J.N.P.). Additional support was provided by Achievement Rewards for College Scientists (ARCS) and a National Cancer Institute NRSA F31 fellowship F31CA284712 (to D.S.B).

## Author contributions

**D.S.B:** Conceptualization, Investigation, Methodology, Validation, Writing - Original Draft, Visualization.

**R.E.L:** Investigation, Resources, Writing - Review & Editing.

**J.N.P:** Conceptualization, Resources, Writing - Review & Editing, Funding acquisition.

**M.S.C:** Conceptualization, Supervision, Writing - Review & Editing, Funding acquisition.

