## Supplemental Materials for "Discovery of ester-linked ubiquitylation of PARP10 mono-ADP-ribosylation in cells: a dual post-translational modification on Glu/Asp side chains"

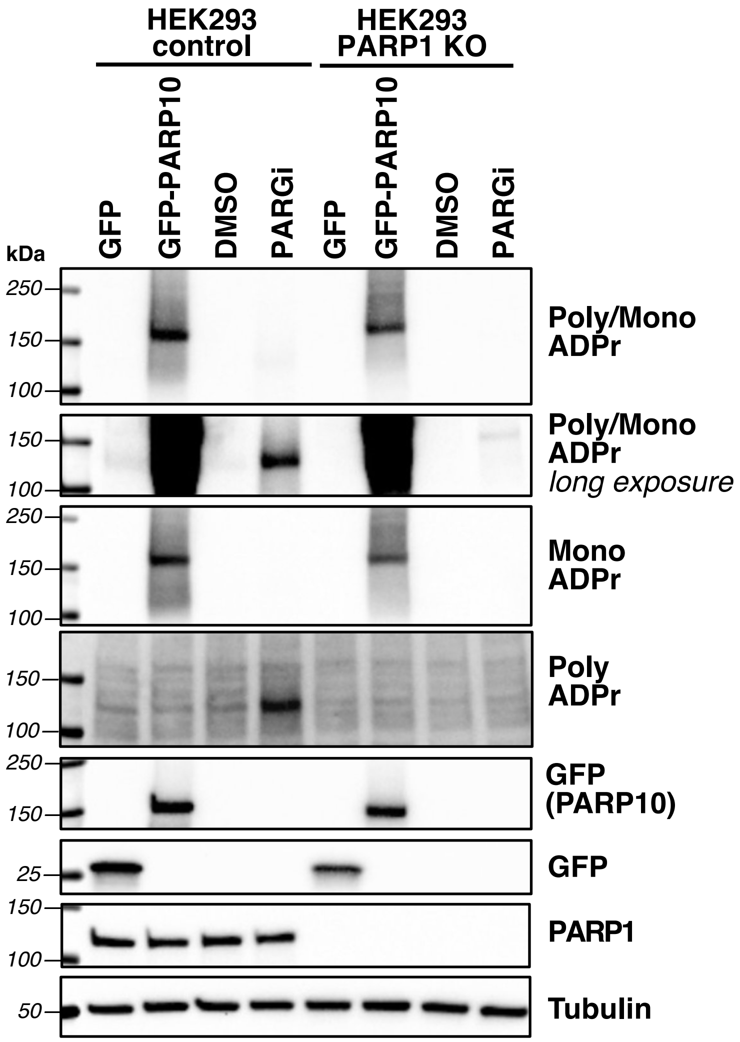


**Figure S1. Evaluating specificity of poly-ADPr, mono-ADPr, and poly/mono-ADPr antibodies.** HEK 293 control and PARP1 KO cells were transiently transfected with GFP or GFP-PARP10 for 24 hours, or treated with PARG inhibitor (PDD00017273, 1 μM) for 30 minutes, followed by western blotting and probing for poly/mono ADPr (Cell Signaling Technology: E6F6A), mono-ADPr (Bio-Rad: HCA354), or poly-ADPr (Millipore Sigma: MABE1031).

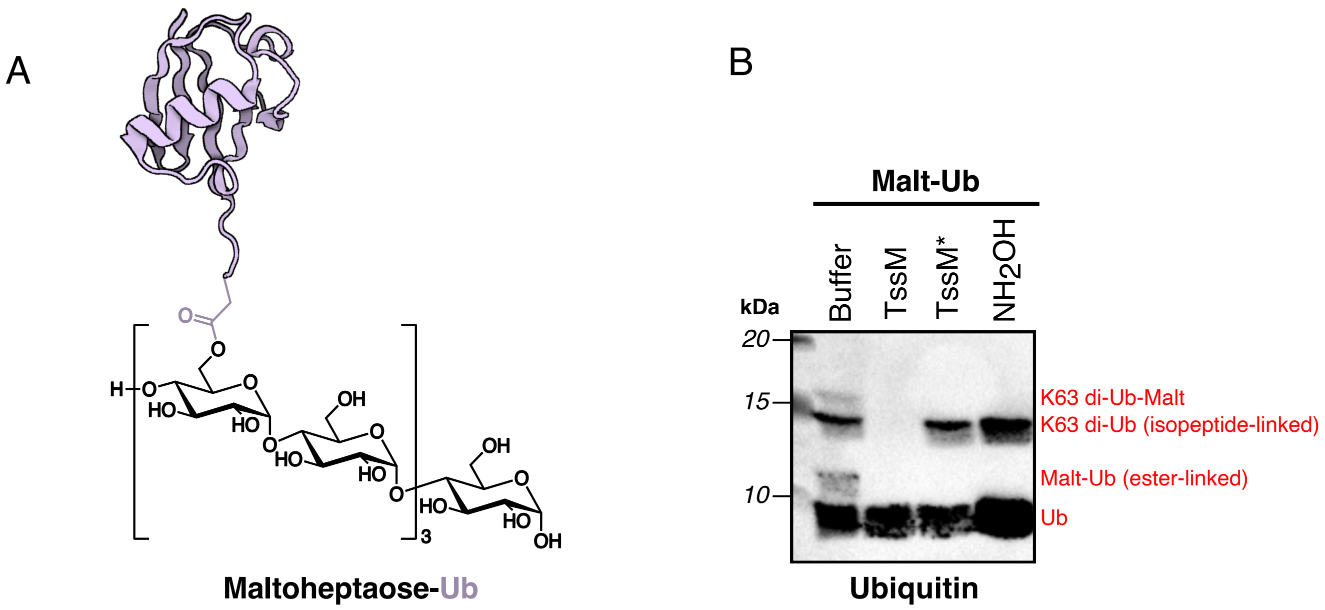


**Figure S2. TssM* is highly specific for removal of ester-linked maltoheptaose-Ub.**

(A) Structure of maltoheptaose-Ub (Malt-Ub) with an ester-linked Ub on the C6 hydroxyl group of glucose. (B) Malt-Ub, generated as previously described (Szczesna et al 2024), was incubated with 2 μM TssM, 2 μM TssM* or 1M $NH_2OH$ (pH 7.5) for 1 hr at 37ºC.

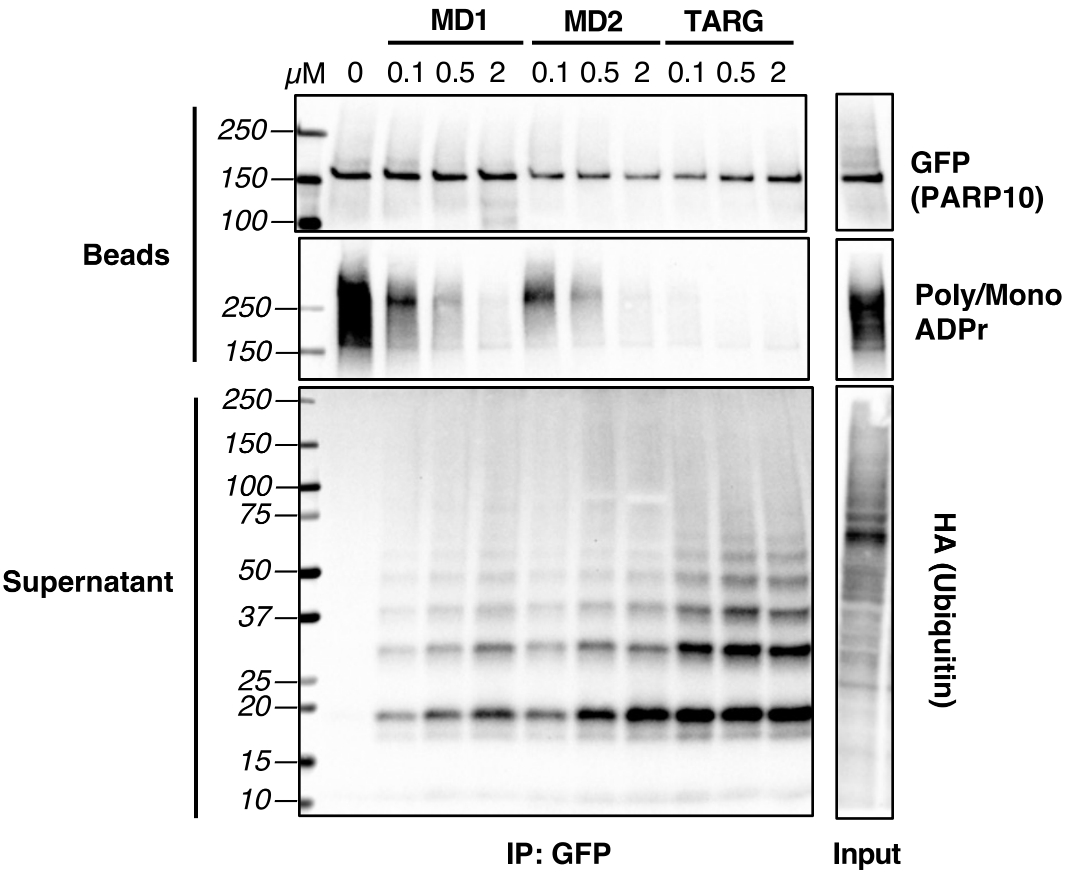


**Figure S3. Screening macrodomain-based ADPr hydrolases against PARP10 MARylation**

GFP-PARP10, from doxycycline-induced HEK 293 cells transfected with HA-Ub, was immunoprecipitated with GFP-trap beads, washed stringently (7M Urea, 1% SDS), and beads were treated with a dose response of MacroD1 (MD1), MacroD2 (MD2) and terminal ADP-Ribose protein glycohydrolase (TARG). The supernatant and bead fractions were separated and subjected to western blotting. Representative image from n = 2 biological replicates.

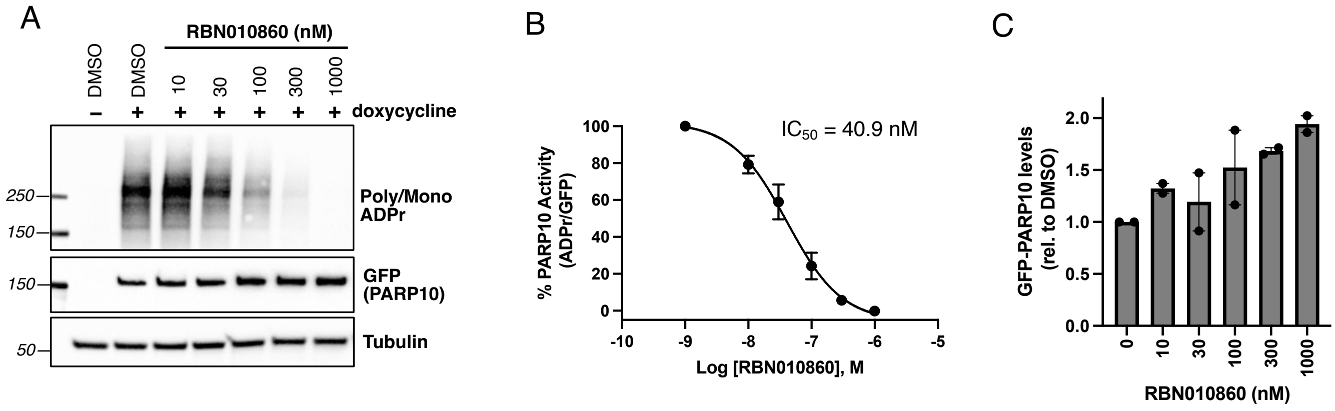


**Figure S4. RBN010860 potently inhibits PARP10 MARylation and leads to dose-dependent increases in PARP10 levels.**

(A) HEK 293 GFP-PARP10 doxycycline-inducible cells were co-treated with doxycycline (10 µg/ml) and a dose response of RBN010860 for 24 hours, followed by western blotting. Quantification of the $IC_{50}$ value of RBN010860 (B) and concomitant increase GFP-PARP10 levels (C); n = 2 biological replicates.

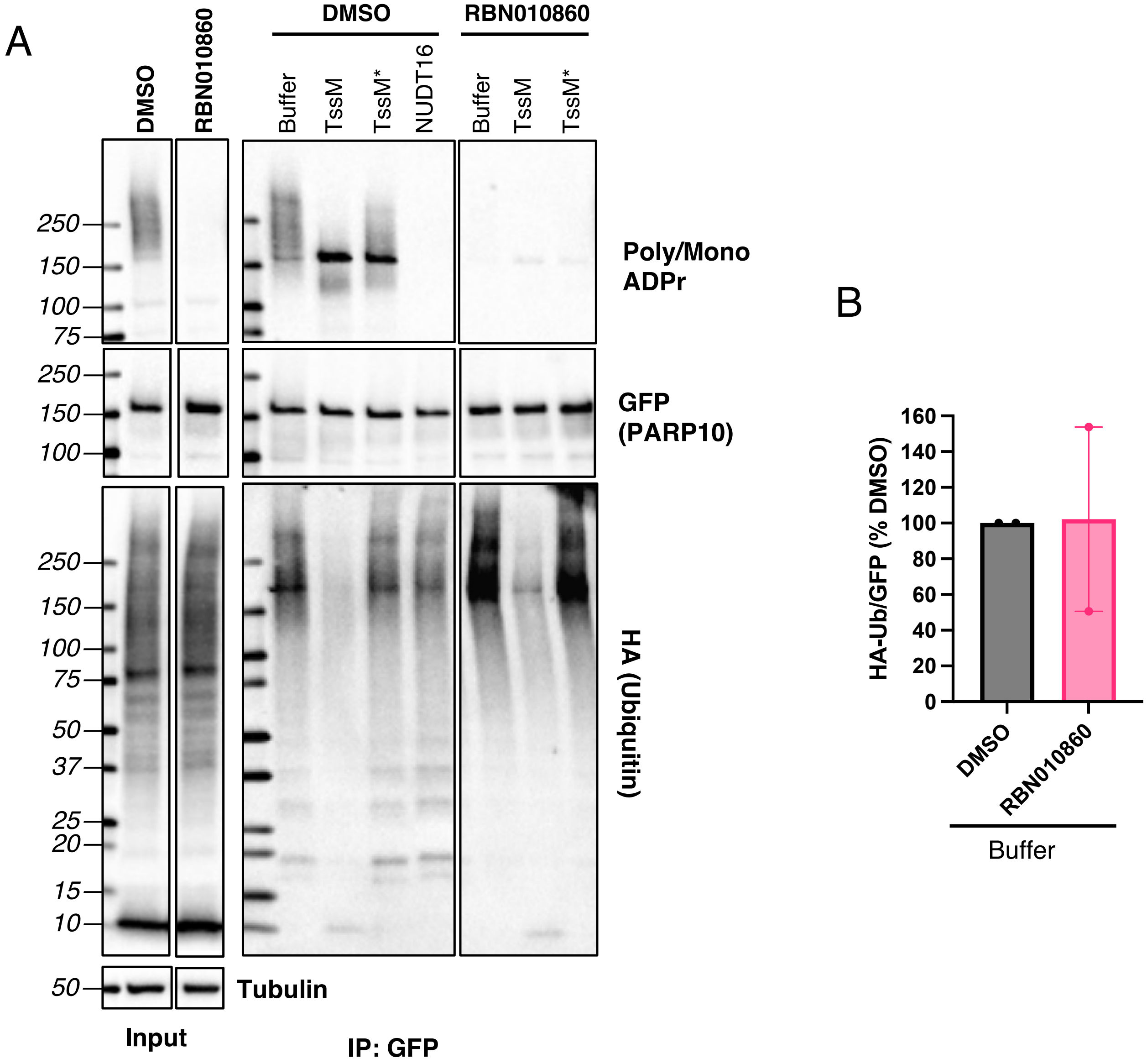


**Figure S5. PARP10 contains a population of isopeptide-linked polyUb that is not dependent on MARylation.**

(A) GFP-PARP10 dox-inducible HEK 293 cells were transfected with HA-Ub and treated with DMSO or 1 μM RBN010860 for 24 hr, followed by immunoprecipitation with GFP-trap beads, stringent washing (7M Urea, 1% SDS), and on-bead treatment with TssM (1 μM) or NUDT16 (10 μM). (B) Quantification of the HA-Ub signal (normalized to GFP) from buffer treated samples from DMSO or RBN010860 treatment; n = 2 biological replicates.
